# Neural Activity Shaping in Visual Prostheses with Deep Learning

**DOI:** 10.1101/2023.12.20.572123

**Authors:** Domingos Castro, David B. Grayden, Hamish Meffin, Martin Spencer

## Abstract

**Objective:** The visual perception provided by retinal prostheses is poor and limited to images constructed of phosphenes generated by the electrodes. One limiting factor has been the conventional strategy used to encode the target image into a stimulation pattern. Under this strategy, if the electrode density is high, the current spread of neighbouring unipolar stimuli overlaps, leading to blurred images. Simultaneous multipolar stimulation guided by the measured neural responses can attenuate excessive spread of excitation and allows for a more precise electrical input to the retina. However, it is far from trivial to predict what multipolar stimulus pattern will elicit the desired retinal response for a given target image. Here, we propose to solve this problem using an Artificial Neural Network (ANN) that could be trained with data acquired from the implant itself.

**Approach:** Our method consists of two ANNs trained sequentially. The Measurement Predictor Network (MPN) is trained on data from the implant and is used to predict how the retina responds to multipolar stimulation by learning the forward model. The Stimulus Generator Network (STG) is trained on a large dataset of natural images and uses the trained MPN to determine efficient multipolar stimulus patterns by learning the inverse model. We validate our method *in silico* using a realistic model of retinal response to multipolar stimulation.

**Main Results:** We show that the simulated retinal activations elicited with our ANN-based approach are considerably sharper when compared with the conventional method used in existing devices. The SGN finds multipolar stimulation patterns that are tuned to a specific retina, thus providing patient-specific stimuli. Also, due to its small computational cost, the SGN can output stimulation patterns at a very high rate.

**Significance:** Our novel protocol opens the door to personalized multipolar retinal stimulation, which may improve the visual experience and quality of life of retinal prosthesis users.

## 1. Introduction

Vision loss affects approximately 230 million people worldwide (Wang et al., 2023). Electrical stimulation of the retina or cortex have emerged as promising approaches to restore some visual perception for people with severely impaired vision. In diseases such as age-related macular degeneration or retinitis pigmentosa, injecting current into the retina allows for the direct stimulation of the surviving inner retinal cells, bypassing the deteriorated outer retina, where the photoreceptors lie (Humayun, 1996; Weiland et al., 2005). On the other hand, injecting current into the visual cortex bypasses the initial visual pathway allowing for treatment of conditions that damage the visual nerve, such as glaucoma (Brindley & Lewin, 1968; Lewis et al., 2015; Niketeghad & Pouratian, 2019). Visual prostheses typically comprise a camera that captures the image meant to be seen by the patient, an algorithm that transforms this image into a stimulation pattern and a multi-electrode array (MEA) that delivers the stimuli at the specified locations (Figure 1.A). Since visual perception is correlated with the spatial activation of the neuronal cells (be it at the retina or cortex), the goal of existing retinal implants is to activate neurons that encode for bright regions of the image. Throughout this manuscript, we focus on retinal prostheses but an analogous approach might be adapted to cortical implants.

**Figure 1.**
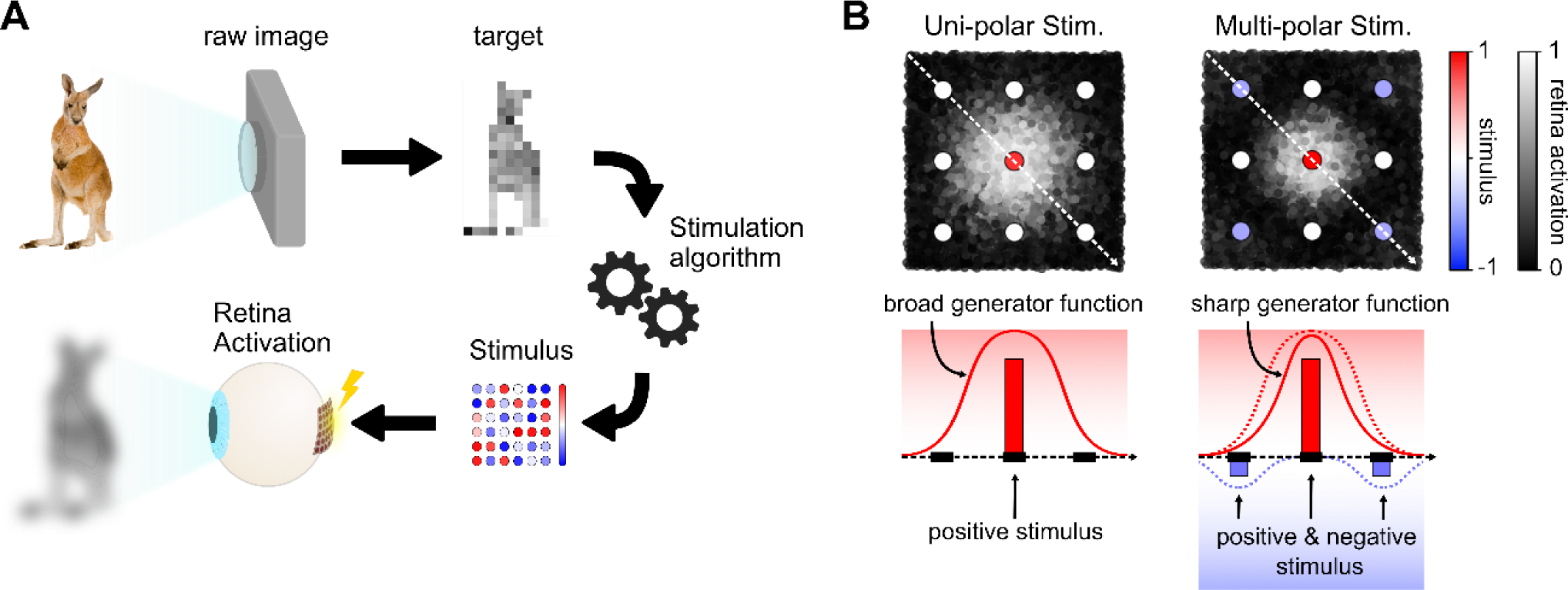
Restoring vision with retina electrical stimulation. A) Schematic representation of the stimulation workflow: a camera captures the desired field of view of the patient; the raw image is subsampled to create a target for the stimulation algorithm; the generated stimulus pattern activates the retina so that the elicited visual perception resembles the original image. B) Sequential unipolar stimulation (either cathodic-first or anodic-first stimulus in all electrodes) leads to broad retinal activations (left) while simultaneous multipolar stimulation (both cathodic-first and simultaneous anodic-first stimulus in different electrodes) can attenuate unwanted current spreads and generate sharper activations.

A critical part of the visual prosthesis workflow is the algorithm used to create the stimulation patterns. A stimulation algorithm typically consists of two main steps: 1) conversion of the captured image into a simplified low-resolution target image and 2) encoding of the target image into an electrode stimulation pattern that ensures the desired retina activation. Most studies on stimulation algorithms aim to solve the first step, by designing methods that preprocess the original image (higher contrast, edge detection, etc.) or highlight important elements of the image (face detection, foreground/background, etc.) as reviewed by Wang et al. (2023). Nonetheless, the second step – encoding the target image into a stimulation pattern – has great potential to impact the acuity of the visual experience.

Currently, stimulation patterns are formed by directly translating the brightness values of the down-sampled target image into stimulation amplitudes at the MEA. The idea is to recreate the target image by combining individual phosphenes (the visual percept elicited by a stimulus) like pixels in a bitmap. However, this method is limited by two main factors: 1) it does not take into account the heterogeneities of the retinal responses to stimulation, which typically lead to distorted phosphenes (Beyeler et al., 2019), and 2) as electrode density increases, the retinal activation induced by neighbouring electrodes overlap, imposing a limit on the sharpness of the perceived images (Tong et al., 2020). This problem is exacerbated when the stimulation is performed with multiple electrodes simultaneously due to spread of current to the inter-electrode regions (Schmid et al., 2013; Wilke et al., 2011).

The blurring problem due to excessive current spreads could be minimized using stimuli of opposite polarities applied simultaneously, leading to an improved visual experience (Figure 1.B). This multipolar strategy has been tested in cochlear implants allowing for focused stimulation (Shepherd et al., 2017; Smith et al., 2013; van den Honert & Kelsall, 2007). Multipolar stimulation has already been tested in retinal implants *in vivo* and was shown to produce spatially restricted phosphenes (Spencer et al., 2016). To properly shape the electric field, they tuned the stimulation amplitudes according to the measurable current spreads. Spencer et al. (2019) proposed that multipolar stimulation could be tuned according to the actual retinal responses, instead of the measurable stimulation current signal. This multipolar strategy was named Neural Activity Shaping. However, the methods presented so far can be computationally very expensive since the number of possible multipolar stimulus patterns is immense and the methods did not take into account some aspects of the non-linear responses to stimulation of retinal cells and were developed based on a simplified retinal model.

In this work, we develop a stimulation algorithm based on Artificial Neuronal Networks (ANNs) that uses Neural Activity Shaping to generate sharp retinal activations. Our ANN-based stimulation strategy is trained using data obtained from the implant itself, so it finds effective stimulation patterns that are patient-specific. Briefly, the method consists of two independently trained ANNs: a Measurement Predictor Network (MPN) and a Stimulus Generator Network (SGN). The MPN learns the forward model and is thus a decoder of the stimulus pattern. It is trained based on patient data and its goal is to accurately predict the neural response to a given multipolar stimulation pattern; more specifically, it predicts the retina response that is *measured* using the implanted MEA. The SGN learns the inverse problem, allowing it to encode the stimulus patterns for a given target image. Once trained, the SGN outputs the stimuli that lead to the desired retinal activations based on the MPN prediction.

This encoder-decoder framework has been recently introduced in the visual prosthesis context (van Steveninck et al., 2022). However, the decoder used by van Steveninck et al. (2022) was not a predictor of the retina response to stimulation but was instead trained to convert the estimated retina response into a new artificial representation similar to the target image. The approaches followed by others (Granley et al., 2022.; Relic et al., 2022; Wu et al., 2023) were more realistic in the sense that the decoders, be they a neuronal network or other differentiable model, were estimators of actual visual perception. However, there are significant distinctions between their approach and ours. In these studies, the goal of the encoders was to define stimulation patterns that accounted for phosphene distortions, characteristic of epiretinal implants, using multi-electrode *monopolar* stimulation. Our encoder, on the other hand, is trained to use Neural Activity Shaping (multipolar stimulation tuned according to the measured retina response) to solve the blurring problem that occurs when the phosphenes overlap. Irregular phosphene shapes are thus learned implicitly. Also, in a clinical context, the decoders presented in the previous studies would estimate an intermediate mathematical model that approximated the phosphenes elicited by stimulation. Their model was parametrized according to the patient’s description of the phosphenes. Imprecisions in this intermediate model, either due to the patient’s description or due to the limited flexibility of the mathematical model, leak into the estimated decoder, which then propagate to the encoder that generates the stimulus patterns. Another possible source of imprecision is that, since the original mathematical model described the phosphenes generated with single electrode stimulation (Beyeler et al., 2019), it did not account for crosstalk between electrodes that may occur when using multielectrode stimulation (Schmid et al., 2013; Tong et al., 2020; Wilke et al., 2011). Our decoder network, on the other hand, is trained directly on the measured retinal responses elicited by the multipolar stimulation. This also allows for online retraining of the encoder network to account for long-term changes, in a closed-loop fashion. Finally, considering the measured responses to stimulation gives us a testing error that is actually measurable in a real scenario, which would not be feasible in the other proposed frameworks.

We test our algorithm using a realistic mathematical model of retinal response to multipolar stimulation presented in our previous *in vitro* study (Maturana et al., 2016). We show that our ANN-based stimulation protocol finds efficient stimulation patterns by combining positive and negative currents. Also, the trained SGN can output the stimulus patterns at a very high rate (much higher than that required for a visual implant). Finally, the stimulation results are significantly better than those obtained with the conventional approach used in existing devices.

Artificial neuronal networks are considered general function approximators. This means that our protocol should lead to an effective and personalized stimulation algorithm regardless of the type of MEA architecture or electrode density. We believe that our strategy will greatly benefit future retinal implants by leveraging on the increased electrode densities – and subsequent stimulation overlap – of high electrode density devices.

## 2. Methods

### 2.1 ANN-based stimulation protocol

The proposed data-driven stimulation protocol requires training two ANNs sequentially (Figure 2). The first network to be trained is the decoder: the Measurement Prediction Network (MPN). The goal of the MPN is to predict how a specific patient’s retina responds to a given multipolar stimulus provided by the implanted MEA. In a clinical context training, the MPN would require a dataset composed of multiple stimulus patterns and the associated retinal responses as training data(see section 2.2 for details). This dataset would be built by submitting a set of random stimuli to the patient using the implant and recording retinal responses that are measured directly by the implant. This stage would take less than 20 minutes, using 10,000 stimuli at a frame rate of 10 stimuli per second. Here, we simulate this step using a computational model of the retina (see section 2.4 for details on the model). The MPN can then be trained offline.

**Figure 2.**
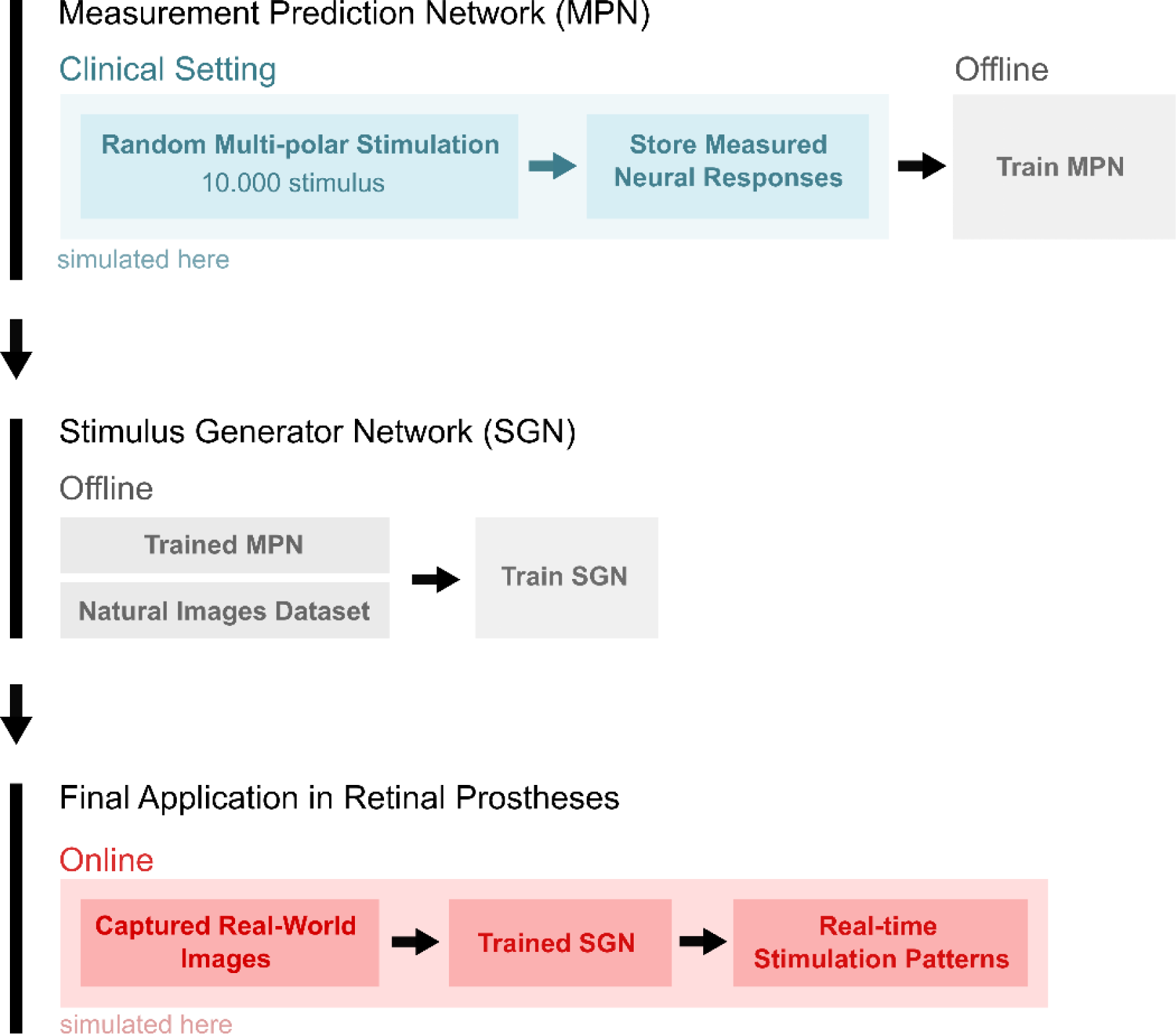
Proposed ANN-based stimulation protocol for visual prostheses. The protocol has three stages: 1) training the MPN based on data of the patient’s retinal response to random stimuli; 2) training the SGN using the trained MPN and a large dataset of natural images; 3) implementing the trained SGN in the implant for real-time generation of stimulus patterns.

Once the MPN is trained (i.e., once it can accurately predict how the patient’s retina responds to a given stimulus pattern), it can be used to train the encoder: the Stimulus Generator Network (SGN). This network outputs the multipolar stimulus pattern that elicits the desired retinal response for a given target image. It can be trained purely offline using a large dataset of natural images and the MPN, which is used to evaluate how similar the evoked retinal response is to the target image (see section 2.3 for details).

Once the SGN is trained, its parameters can be integrated into the implant to run in real time. The images captured with a camera would be pre-processed (not in the scope of this work) and sent to the SGN, which would produce effective multipolar stimulation patterns. This stage is also simulated in our work using the same retina model.

It is important to note that, even though we are testing this protocol in a simulated environment (where we as developers can access all the variables of the simulation), all the methods presented in this work were designed to be dependent only on the information accessible through the implant; i.e., the delivered stimulus patterns and the measured retinal responses. This ensures that the algorithms can be directly applied in a clinical setting. Also, this framework is not limited to retinal implants; it mayalso be adapted for cortical visual protheses, so long as the transformation from target image to cortex representation is known in order to create the target activity patterns.

### 2.2 Measurement Predictor Network

The MPN is the data-driven, patient-specific predictor of the retinal responses to stimulus (Figure 3). It is a fully connected ANN having the stimulus pattern as input, two hidden layers, and the predicted measured retina response as output layer. In this study, we consider a square MEA with an 8-by-8 matrix of electrodes. Thus, the input and output layers are vectors with 64 neurons. The two hidden layers have 128 neurons each. We used the Rectified Linear Unit (ReLU) activation function in the hidden and output layers. This architecture was optimized empirically with the aim of obtaining a small but effective network. We used the open-source Keras API to build and train the ANNs, running on a i7-1165G7 processor at 2.80 GHz.

**Figure 3.**
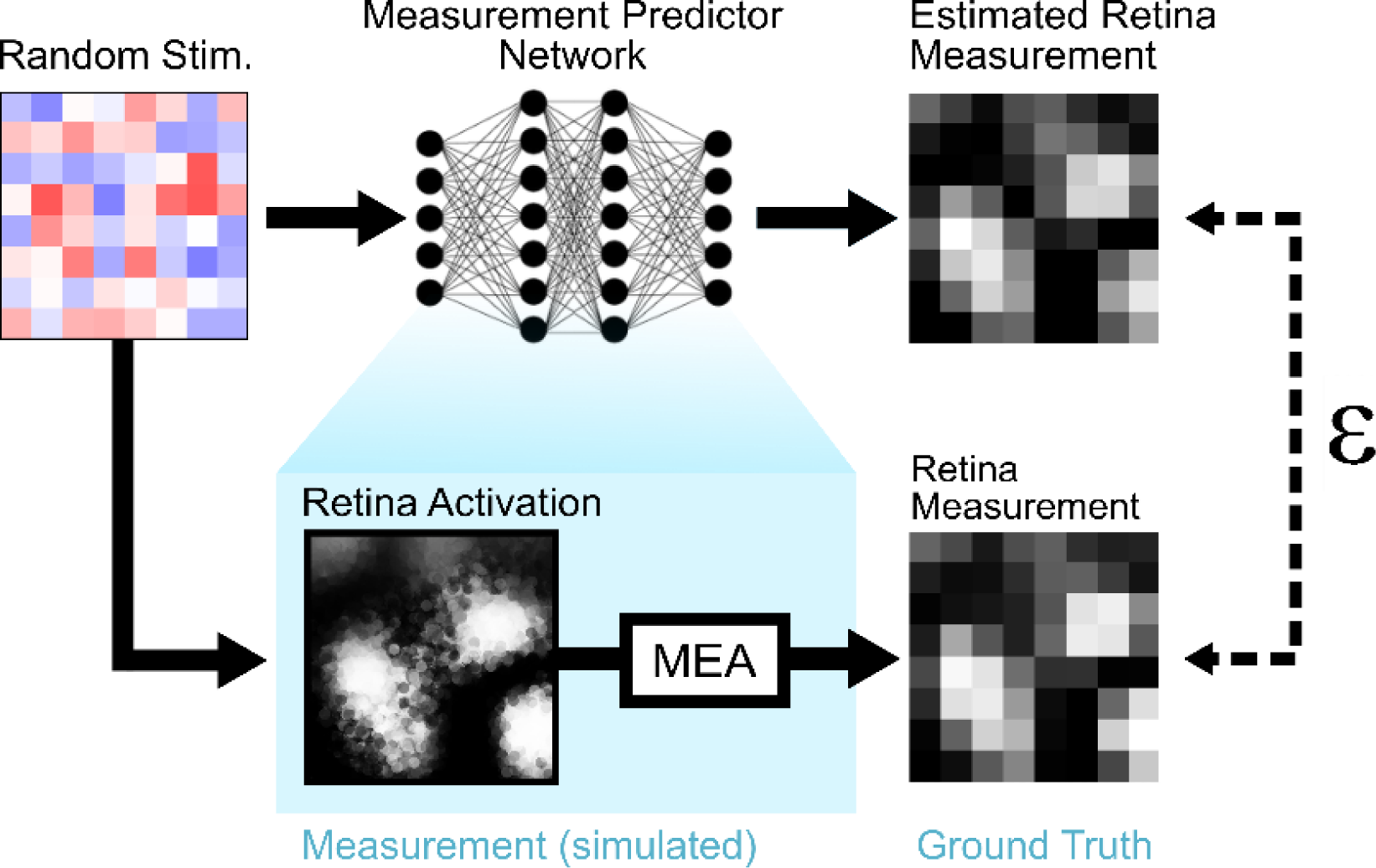
Training of the Measurement Predictor Network. The MPN is a decoder that predicts the measurement of the retinal response to a given stimulus. It is trained with a dataset composed of multiple stimulus patterns and the associated measured retinal response. Here, both the stimulation and the recording are performed with the implanted MEA (simulated in this study) but could also be done with separate stimulating and recording units.

The MPN is trained using random stimulation patterns. The amplitudes of the multipolar stimuli are randomly sampled from a Gaussian distribution with mean 0 and standard deviation 0.3 to limit the stimulation amplitudes between –1 and +1. The stimulation units considered in this study are arbitrary, but this has no implication on the applicability of the MPN in a real clinical setting. The ground truths are the measured retinal response to stimulation. In a clinical setting, these are obtained by recording the retinal activation after each stimulus, requiring the use of an implant with a neural recording ability. Here, we simulate the retinal response to stimulus and the recording performed with the MEA (see section 2.4 for details). Just as in a clinical setting, the ground truths in our simulated environment are the recorded responses.

The split between training, validation, and testing sets are 10,000, 2,000 and 1,000 stimulus patterns, respectively. We use the ADAM optimizer with an initial learning rate of 0.0005 (the remaining parameters were set to their default value). As loss function, we use the Mean Square Error between the retinal response predicted by the MPN and the actual retinal response measured with the MEA (Figure 3).

To evaluate the robustness of the MPN, we train it using different initial conditions by changing both the seeds of the random number generator and retinal activation spread. The latter is a parameter that controls the spatial extension of the retinal activation for a given stimulus (see details in section 2.4).

### 2.3 Stimulus Generator Network

The SGN receives a target image and outputs a patient-specific multipolar stimulation pattern. The target image provided to the SGN has the resolution of the MEA because it needs to be compared with the measured retinal response or its prediction during training (Figure 4). We refer to this down-sampled target as the *measurable target* to distinguish from the original target image captured with the camera. Here, the measurable targets are created by filtering the original images with a low-pass filter to avoid aliasing and down-sampling it to the MEA resolution.

**Figure 4.**
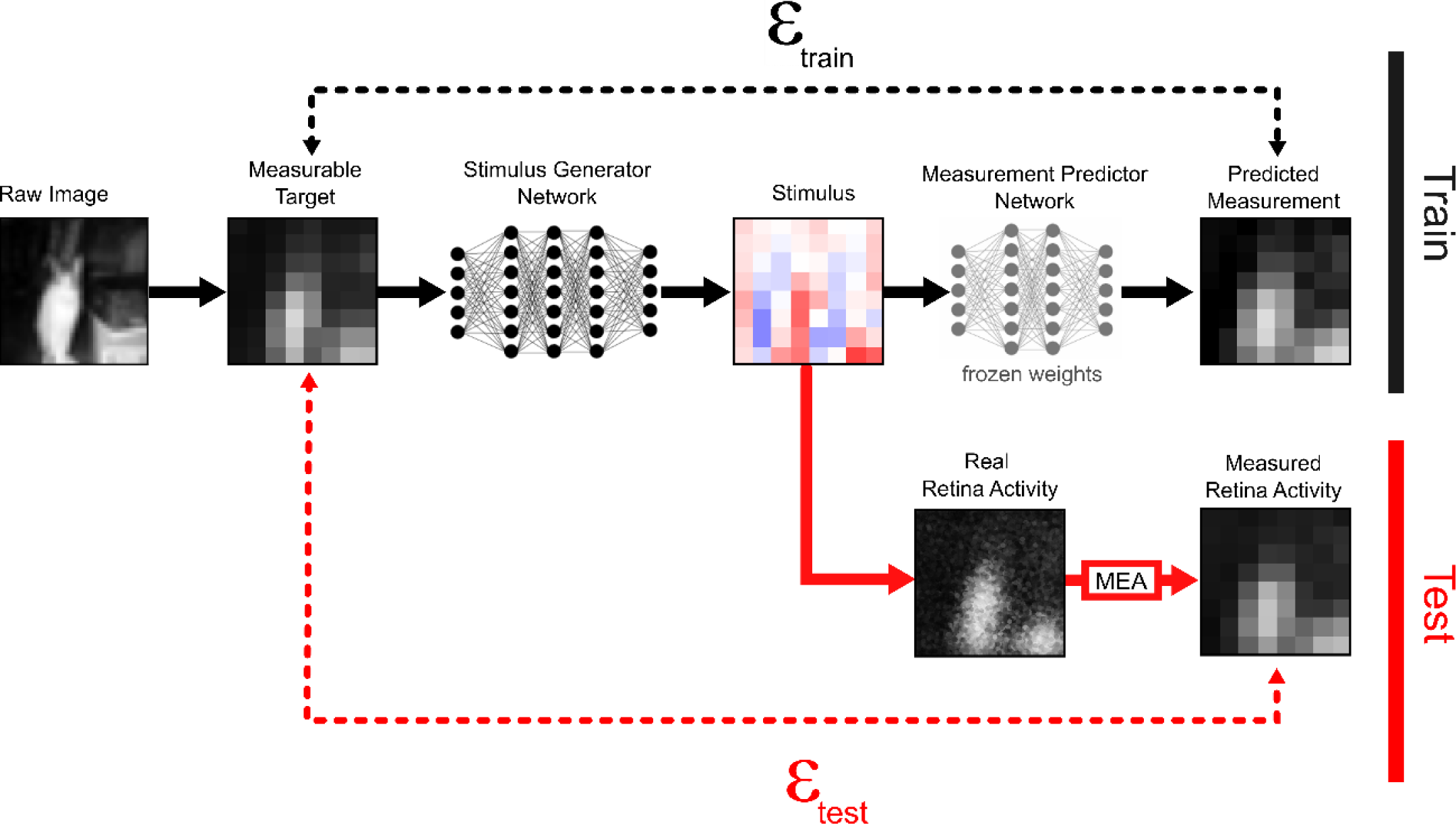
Training and testing of the Stimulus Generator Network. The SGN receives a subsampled version of the raw captured image as input – the Measurable Target; this is the desired retinal response that we want to measure with the MEAs. The SGN outputs a stimulation pattern that should elicit the desired retinal response. To train the SGN, we use the patient-specific predictor of retinal response, the MPN, and a large dataset of natural images (black branch). Once the SGN is trained, it can be tested in the real retinal implant (red branch, here simulated).

The SGN has three hidden layers, each with 128 neurons and ReLU activation functions. The values of output layer correspond to the stimulation amplitudes of each electrode of the MEA, thus having 64 neurons. The output layer has a hyperbolic tangent activation function to bound the stimulation amplitudes between –1 and +1. The architecture of the network is optimized empirically to create a small but effective network (optimization results not shown). Finally, we add L2 regularization to the output layer to penalize high amplitude stimulus. We tested the following values for the *lambda* parameter of the L2 regularization: 0, 10^−5^, 10^−4^, 10^−3^, 10^−2^ and 10^−1^.

To train the SGN, we merge it with the trained MPN. The weights of the MPN are frozen so that the only trainable parameters are those from the SGN. Stacking these two networks allows the training of the SGN to be performed offline using a large dataset of natural images since the MPN predicts how the patient’s retina responds to the stimulus outputted by the SGN (black branch in Figure 4). For this study, we use the CIFAR 100 dataset, which contains 60,000 32-by-32 images (Krizhevsky, 2009). The split between training, validation, and testing sets is 37,500, 12,500, and 10,000 images, respectively. The training is performed for 300 epochs using the ADAM optimizer with an initial learning rate of 0.0005. As the loss function, we use the mean-squared error between the measurable target and the predicted measurement outputted by the MPN. After the training is completed, the trained SGN is tested on the actual retina (here simulated) and the measured retinal responses are compared with the test set of measurable targets (red branch in Figure 4).

### 2.4 Model of Retina Stimulation and Recording

The proposed stimulation protocol is tested in a simulated environment composed of *N*r retinal cells evenly distributed in a 2D surface, overlaid with a matrix of *N*e stimulating/recording electrodes. In our simulations, we consider *N*r = 10,000 and *N*e = 64. The cells are placed on a square grid and randomly jittered to create a uniform, yet random, retina. For mathematical convenience, the grid has dimensions 32-by-32 (unitless) to match the 32-by-32 image dataset. The response rate of the retina to electrical stimuli, 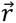, is simulated using a previously presented mathematical model that describes the response of individual retinal cells to simultaneous multipolar electrical stimulation (Maturana et al., 2016). The model is composed of a linear and a non-linear part (Equation 1). The linear part is referred to as the Generator Potential, 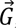, and may be interpreted biophysically as relating to the current crossing the membrane of a given retinal cell as a result of the linear sum of the multi-electrode stimulation (Esler et al., 2018). The contribution of each electrode to the activation of a neuron is given by its electrical receptive field (ERF). The ERFs of all retinal cells are stored as rows of the matrix ***W*** with dimensions *N*r *× N*e. The Generator Potential, 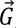, of the entire retina is then given by the dot product of ***W*** with the vector of applied electrode current magnitudes, 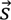.

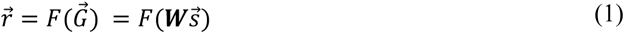

The values of 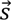 range between –1 and +1, representing maximal cathodic and anodic electrode stimulation, respectively. The ERFs are modelled as 2D ellipsoidal Gaussians. The ERF of each cell *i* has a different angle, *θ*[*i*], and different standard deviations, *σ*_x_[*i*] and *σ*_y_[*i*], for the two axes of the ellipsoids (Figure 5.A). The ellipsoidal ERFs are compiled in ***W*** according to

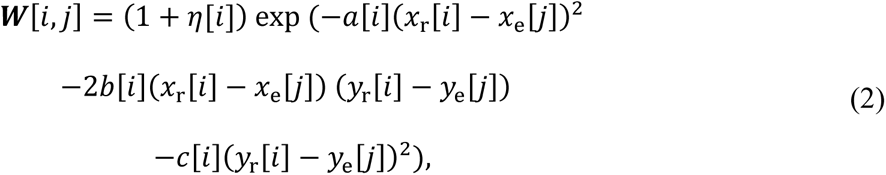

**Figure 5.**
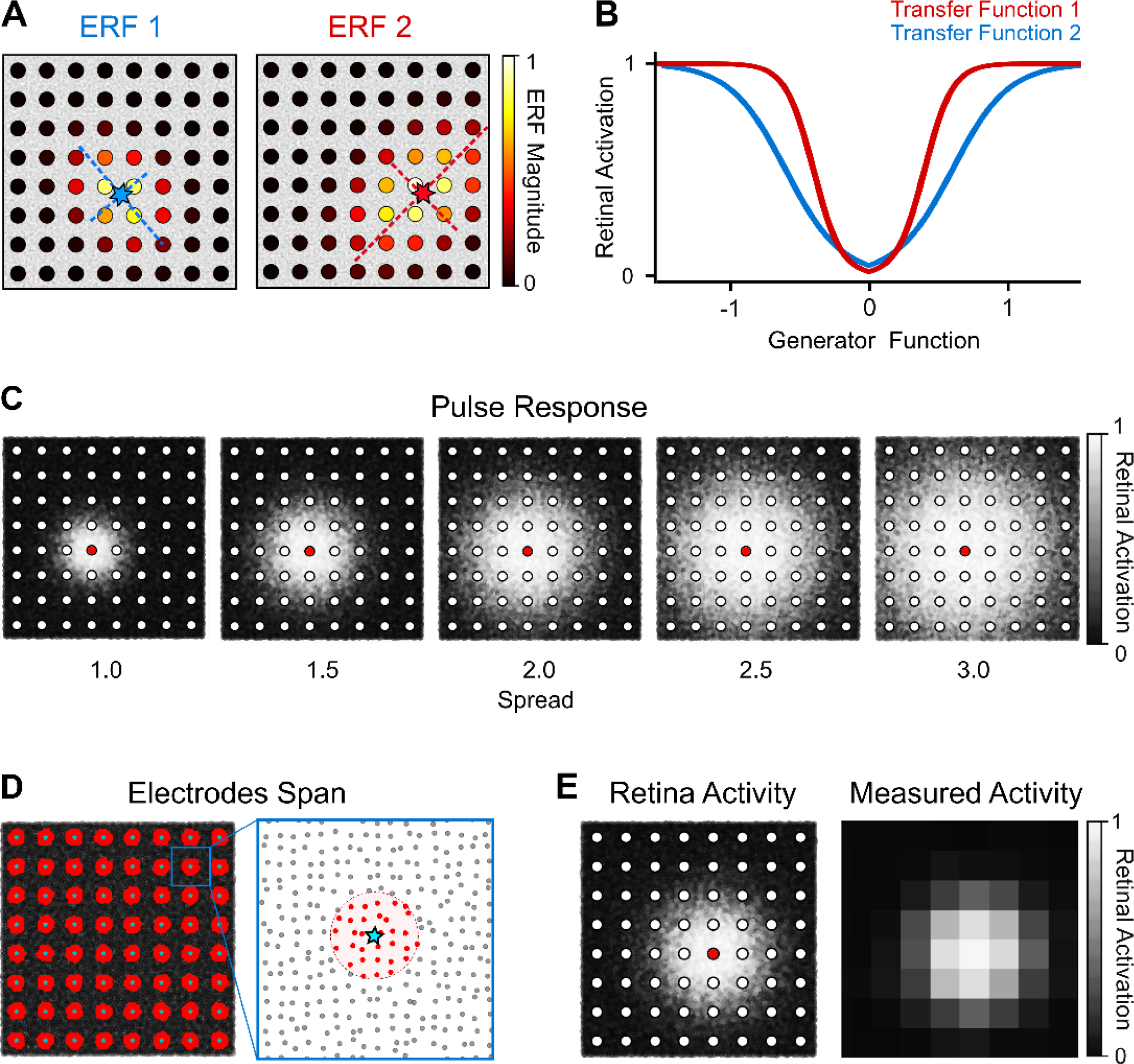
Retina model and MEA recording. A) Representative depiction of the ERFs of two different retinal cells (blue and red stars). B) Representative depiction of two different transfer functions (*k*_1_=5, *z*_1_ = 0.6; *k*_2_=10, *z*_2_ = 0.4). C) Retinal responses to a stimulus of amplitude 1 on the red electrode, for the different spread regimes considered in this study. D) Retinal cells (red) recorded by each electrode (blue star). E) Retina activation that results from a stimulus of amplitude 1 applied in the red electrode (left) and the resulting MEA read-out (right).

where *W*[*i*, *j*] is a weighting factor describing the influence of current from the electrode *j* on the retinal cell *i*, *η*[*i*] is a factor relating the amplitude of response of neuron *i*, here a random number uniformly distributed between –1 and 1, to account for heterogeneities of the retina. *x*_r_[*i*] and *y*_r_[*i*] are the coordinates of the cell *i* and *x*_e_[*j*] and *y*_e_[*j*] are the coordinates of electrode *j*. The values of *a*, *b*, and *c* are

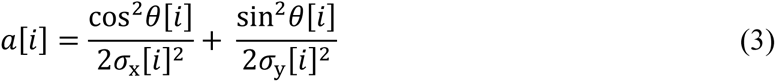

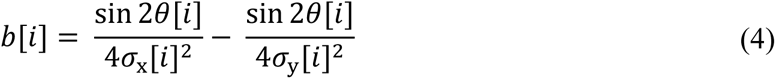

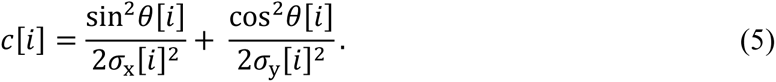

The values of *σ*_x_[*i*] and *σ*_y_[*i*] determine the spatial extension of the ERFs. Each *σ* is obtained using

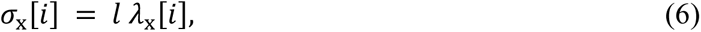

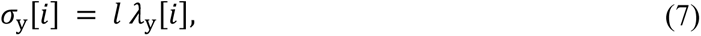

where *l* is the distance between electrodes and *λ* is a dimensionless parameter that determines the extension of the ERF with regards to the electrodes. Large values of *λ* create large ERFs, meaning that the retinal cells are affected by stimuli that are several electrodes away. The values of *λ*_x_[*i*] and *λ*_y_[*i*] for each cell *i* are uniformly sampled from the interval *Λ* ± 25% where *Λ*, also referred to as *spread*, modulates the current overlap of neighbouring electrodes. It can also be seen as a parameter that scales the dimensions of the MEA with regards to the retina (a high-density MEA would be in a high spread regime, for example). We perform simulations under different spread regimes, with *Λ* set to 1.0, 1.5, 2.0, 2.5, and 3.0 electrode pitches.

The non-linear part of the model converts the Generator Potential into retinal activation. This non-linear Transfer Function can be simplified to a double sigmoid function, as previously described in our *in vitro* study (Maturana et al., 2016). Here, we assume that the double sigmoid is symmetric,

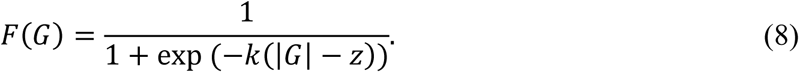

Each cell has a different Transfer Function (Figure 5.B), obtained by uniformly sampling *k* and *z* from the intervals [5, 10] and [0.4, 0.6], respectively. Implementing the realistic linear/non-linear model under different spread regimes allows us to evaluate how our stimulation protocol performs in diverse conditions (Figure 5.C). Elongated phosphenes can also be created by forcing *λ*_x_ and *λ*_y_ to be different and imposing a limited range on the angle *θ* (Figure S1).

We also model the recording of the retinal activity with the MEAs. For that, we assume that the electrodes have a diameter equal to half the electrode pitch (centre to centre) and that each electrode’s read-out is proportional to the average activity of all the cells within its radius (Figure 5.D and E).

## 3. Results

### 3.1 The Measurement Predictor Network accurately estimates retinal responses to simulated multipolar stimulation

The first step of our stimulation protocol consists of obtaining a patient-specific predictor of the retinal response to multipolar stimulation (section 2.1). To train the MPN, the simulated retina is used to calculate the response to 10,000 random stimuli.. We trained MPNs under different spread regimes. To check the consistency of the results for each spread, we replicated each condition five times, each with a different parametrization of the MPN and retinal model (the random number generator initialized with a different seed for each repetition). The training converged smoothly for all conditions.

Once trained, we submitted the MPNs to 1,000 test stimuli. We used the Root Mean Square Error (RMSE) as a metric to evaluate the performance of the network. The RMSE quantifies the *Measurement Error*, the error that could be measured in a clinical setting by comparing the measured retina response with the measurable target. The prediction error was similar for all spreads and consistent across the multiple initializations of the same spread (Figure 6.A). The predicted results were qualitatively very similar to the measurement of the simulated MEA (Figure 6.B and C). This demonstrates that the NNs accurately predicted what will be measured with the MEA by looking only at the stimulus pattern. This is a necessary requirement if the MPS is to be used to train the SGN. The MPN was efficient for all level of spread (Figure 6.A). This is an important observation since this variable cannot be controlled in a real scenario as it is intrinsic to the retinal implant system.

**Figure 6.**
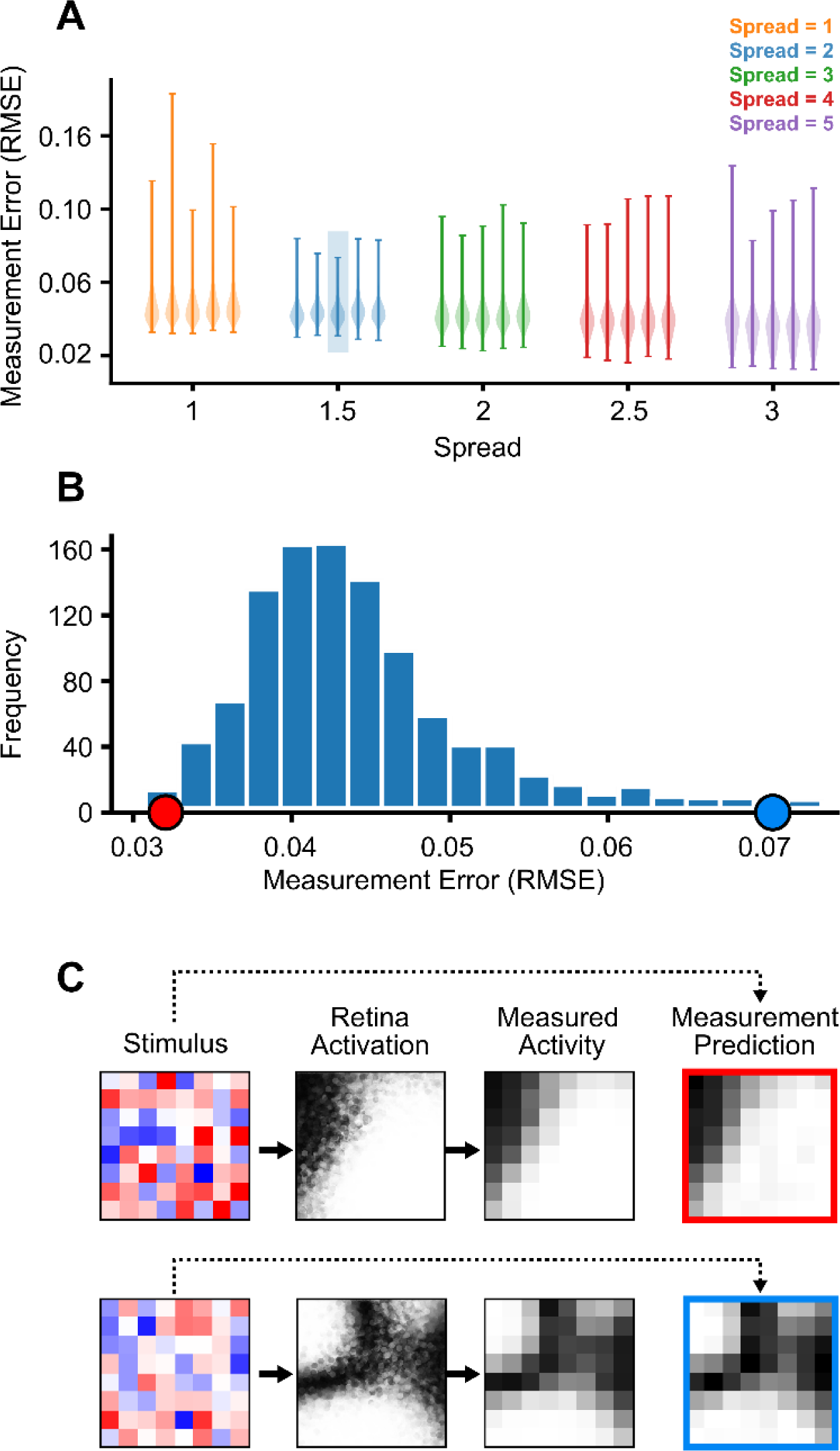
The Measurement Predictor Network predicts the stimulus response in diverse spread conditions. A) Prediction error in the test set for MPNs trained under different spread regimes. For each regime, the experiments were repeated five times with different initializations of the MPN and retina model. B) Histogram of the test error for a spread equal to 1.5 (third repetition, as marked in A). Red and blue marks show the RMSE values of the two qualitative results shown in C. C) Two test stimuli and the associated retina activations, measured activities, and measurement predictions, extracted from the tails of the histogram, showing that the predictions are qualitatively very similar to the actual measurements.

### 3.2 The Stimulus Generator Network produces effective stimulation patterns

The SGN was trained offline using the previously trained MPN and a large dataset of natural images, as explained in section 2.3. Each MPN-SGN pair was trained within the same simulation, meaning that the different spread regimes and initializations used for the MPN were also applied for the SGN. After the training, the SGN was tested in a simulation with 10,000 test images and applying the resulting stimulus using the retina model (the one used to train the MPN). We then compared the measured retinal response with the measurable target (red part in Figure 4). For each condition (spread regime and random initialization), we tested six levels of L2 regularization (Figure 7.A). Applying regularization on the SGN output has two effects: it explicitly forces the network to choose lower stimulation amplitudes, which is important in a clinical application, and it implicitly smooths a loss function, making the training less prone to land in sharp local minimum, which improves the generalization of the network.

**Figure 7.**
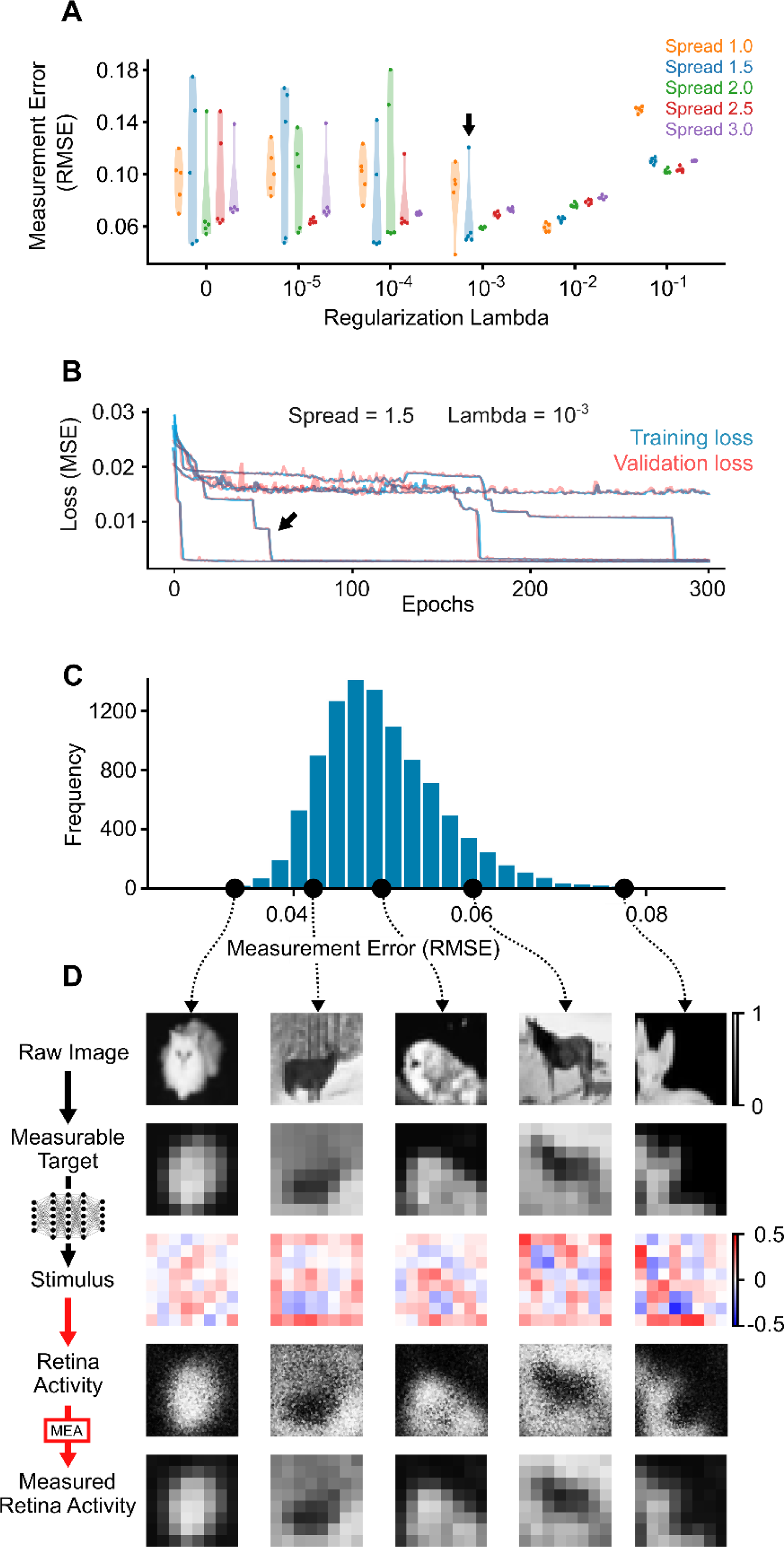
The SGN encodes efficient multipolar stimulation patterns. A) Test results for all the conditions simulated, where each dot represent the average error of 10,000 test stimuli for a given spread, regularization lambda, and random initialization. B) Training loss curves for the five SGNs instantiations at a spread of 1.5 and regularization lambda of 10^−3^ (black arrow in A). C) Histogram of the error for the 10,000 test images for the best SGN instantiation at a spread of 1.5 and lambda of 10^−3^ (whose training is marked with a black arrow in B). D) Qualitative assessment of the results across the spectrum of errors, showing the original raw image, the measurable target that the SGN aims to reproduce, the stimulus pattern produced by the SGN, the retina activation (unattainable in a real scenario), and the measured retinal activity.

Here, we can see that, as the relative magnitude of the regularization increases, the results became more consistent across the multiple initializations for all the spreads. This is due to the smoothing of the loss function. On the other hand, larger regularisation induced lower stimulus amplitudes, which led to less effective stimulation patterns. We found a good compromise between these two effects for a regularization lambda of 10^−3^ (see Methods). Yet, for smaller spreads (1 and 1.5), the SGN may still become stuck in a local minimum even after 300 training epochs (Figure 7.A and B). This means that, in a real application where the actual spread in not known, since it is an intrinsic variable of the retina-implant system, the SGN would need to be trained several times with different initializations and the instance with the best performance would be implemented in the retinal prosthesis. Analogously, here we looked at the results obtained with the best SGN instance for a spread of 1.5 (this spread regime was chosen as an example). The training loss curves show that only one out of five instantiations of the SGN became stuck in a high-error local minimum (Figure 7.B). The best SGN instance for this spread produced results that were qualitatively very similar to the original image for the entire spectrum of test errors (Figure 7.C and D). Finally, it is interesting to note how larger spreads led to better training convergence, which also suggests that the loss function becomes smoother as the overlap between electrodes increases.

The inference time of a trained SGN for 10,000 images running on a i7-1165G7 processor at 2.80 GHz was 766 ± 31 µs (based on 10 test runs, each with 10,000 test images). With this small network, each stimulation pattern was generated in less than 1 µs.

### 3.3 ANN-generated stimulus patterns lead to improved retinal representations compared to the conventional approach

We evaluated how our ANN-based stimulation protocol compared with the conventional stimulation strategy. In the conventional method, the stimulus magnitude of a given electrode is proportional to the brightness of the image in the location underlying the electrode.. The stimulus is typically applied one electrode at a time (sequential stimulation) but fast enough that the patient perceives a single image. Here, we are simplifying this sequential approach by applying the stimulus in all electrodes at the same time and assuming that the visual experience would be identical. For each spread, we tested 10 different scaling factors, ranging from 0.02 to 0.2 in steps of 0.02 (Figure S2). For each spread, we chose the best scaling and compared the stimulation results with those obtained with the best version of the SGN for the same spread.

We first compared the measurable error, which compares the recorded retinal response with the measurable target. For all spreads, the ANN-based approach achieved significantly better results, both quantitatively and qualitatively (Figure 8.A). We also checked the actual retinal activation error; i.e., the error between the actual retinal activity and the original raw image. Since the original image had a 32-by-32 resolution, we approximated the activity of the 10,000 retinal cells to a 32-by-32 image and computed the RMSE between that and the original image (Figure 8.B). The quantitative results showed a comparable improvement to those using the measurable error.

**Figure 8.**
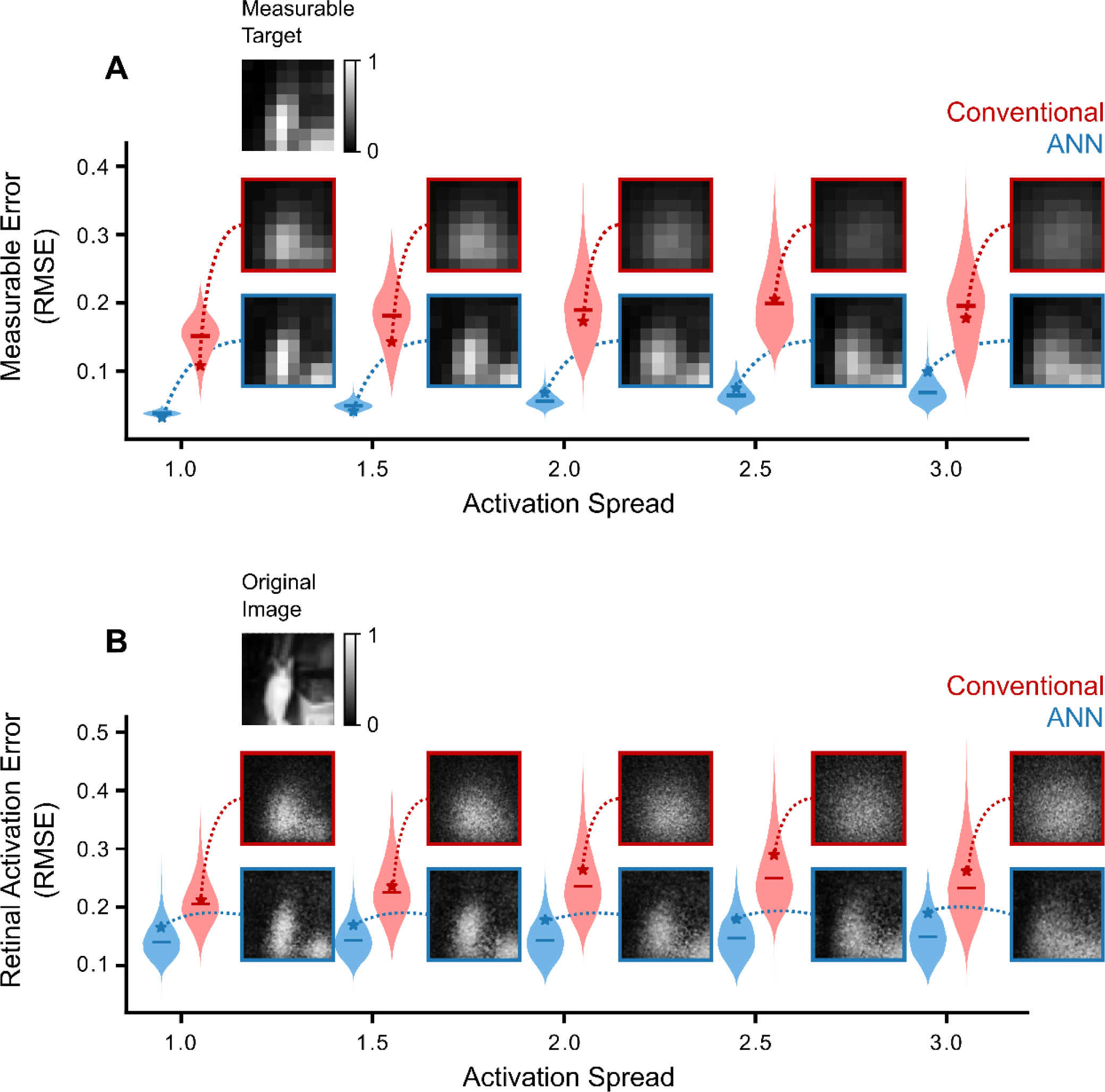
Comparison of conventional and ANN-based stimulation protocols for different spread regimes. A) Measurable error based on the MEA recordings. B) Retinal activation error.

## 4. Discussion

Our ANN-based approach solves the problem of encoding target images into personalized multipolar stimulation patterns. The preprocessing of the captured image into a subsampled target was out of the scope of this work. Here, we simply down-sampled the original image, ensuring no aliasing. But our method could be easily coupled downstream with any of the strategies presented in the literature (Wang et al., 2023) as these are independent tasks. Also, the overall strategy could be applied in any type of device with stimulating and recording capabilities. Still, the specific architecture of the networks used may need to be adjusted; a device with a high-density MEA would generate a more complex dataset and thus require a bigger network, for example.

We tested our approach in a simulated environment, using a realistic model of retinal response to multipolar stimulation. For the sake of simplicity, retinal activation and stimulus magnitude are maxed at 1, but scaling the data to physiological units would have no impact on the results achieved. Despite the heterogeneity of ERFs and transfer functions imposed on the modelled retinas, the visual percepts (phosphenes) elicited were approximately circular, such as those obtained with subretinal implants (R. Wilke et al., 2011). These are also the types of implants that would benefit the most from our strategy because the larger distance between electrodes and ganglion cells leads to considerable current overlap. Likewise, different phosphenes, such as the elongated ones typically seen in epiretinal implants, will impose no problem on the ANNs training, so long as they are fully captured by the recording electrodes (see Fig S1 for examples of results with elongated phosphenes). A limitation of the model considered here is that it is purely spatial and ignores the temporal interactions of the retinal response. We neglect neuronal adaptations such as Long/Short Term Potentiation/Depression, which could change the response to stimulation as a function of time. Another simplification is the assumption that the signal recorded with each electrode corresponds to the average activity of the cells over it. Also, our method does not take into account the fact that the electrical stimulus simultaneously excites cells that may encode different light information (such as ON and OFF ganglion cells), as that is not possible with existing devices. Such specificity would only be possible by recording and stimulating with single cell resolution as well as having the adequate algorithms to identify and target the different cell types (Lotlikar et al., 2023; Madugula et al., 2022; Shah et al., 2019).

One aspect worth exploring in the future is choosing a more suitable error metric for the loss function of the ANNs. Here, we used the standard MSE, but we speculate that metrics sensitive to the information content of the image (such as Structural Image Index) may lead to better outcomes. Also, despite using multipolar stimulus, the final values of the generator functions created with our stimulation method are always unipolar; i.e., the values of 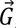 are only positive (or negative) for all the retinal cells and, thus, land only on one side of their Transfer Function. Actually, the SGN instances that become stuck in an inefficient local minimum during training (high error curve in Figure 7.B) were the ones generating bipolar generator functions. When that happens, there are artifacts in the transitions between positive and negative 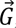 regions (Figure S3). Nonetheless, our previous study showed that it can be highly beneficial to exploit both sides of the non-linearity of the Transfer Function (Spencer et al., 2021) (Figure S3). We suggest that future work could explore how to force the network to use both sides of the non-linearity in order to create higher contrast retinal activations.

Also, we did not show how the number of electrodes affects the stimulation performance (each different M EA configuration would require its own MPN-SGN architectures and parametrizations, which would need to be individually optimized). However, naturally, the higher the number of electrodes, the higher the spatial resolution attainable with the stimulation. Furthermore, as mentioned before, higher electrode density leads to a larger overlap between electrodes, which not only makes our approach more relevant but also more stable, since larger overlaps make the training converge better (even though, for the same number of electrodes, smaller the spreads lead the better image resolution).

In a clinical setting, the networks can be retrained periodically to adapt to the long-term changes of the retinal response. The only requirement is that a subset of the real target images, stimulus patterns and retinal responses are stored in the device. These could be periodically downloaded for retraining the MPN and SGN as before, but now using the actual captured images (instead of random stimuli for the MPN and an arbitrary dataset for the SGN). Another interesting solution would be to do this retraining online, where the weights of the SGN would be changed in real time in a full closed-loop fashion. This way, there would be no need for the MPN during retraining, since the SGN would use the responses measured in real time as ground truths.

To conclude, we believe that our method has great potential to improve the visual experience of retinal prostheses. A critical future step will be to implement our method in an *in vivo* or *in vitro* setup. The significance of this novel approach will increase with the development of higher density electrode implants since, unlike the conventional method, it is not limited by the electrodes overlap, but rather takes advantage of it.

## Supplementary Information

**Figure S1.**
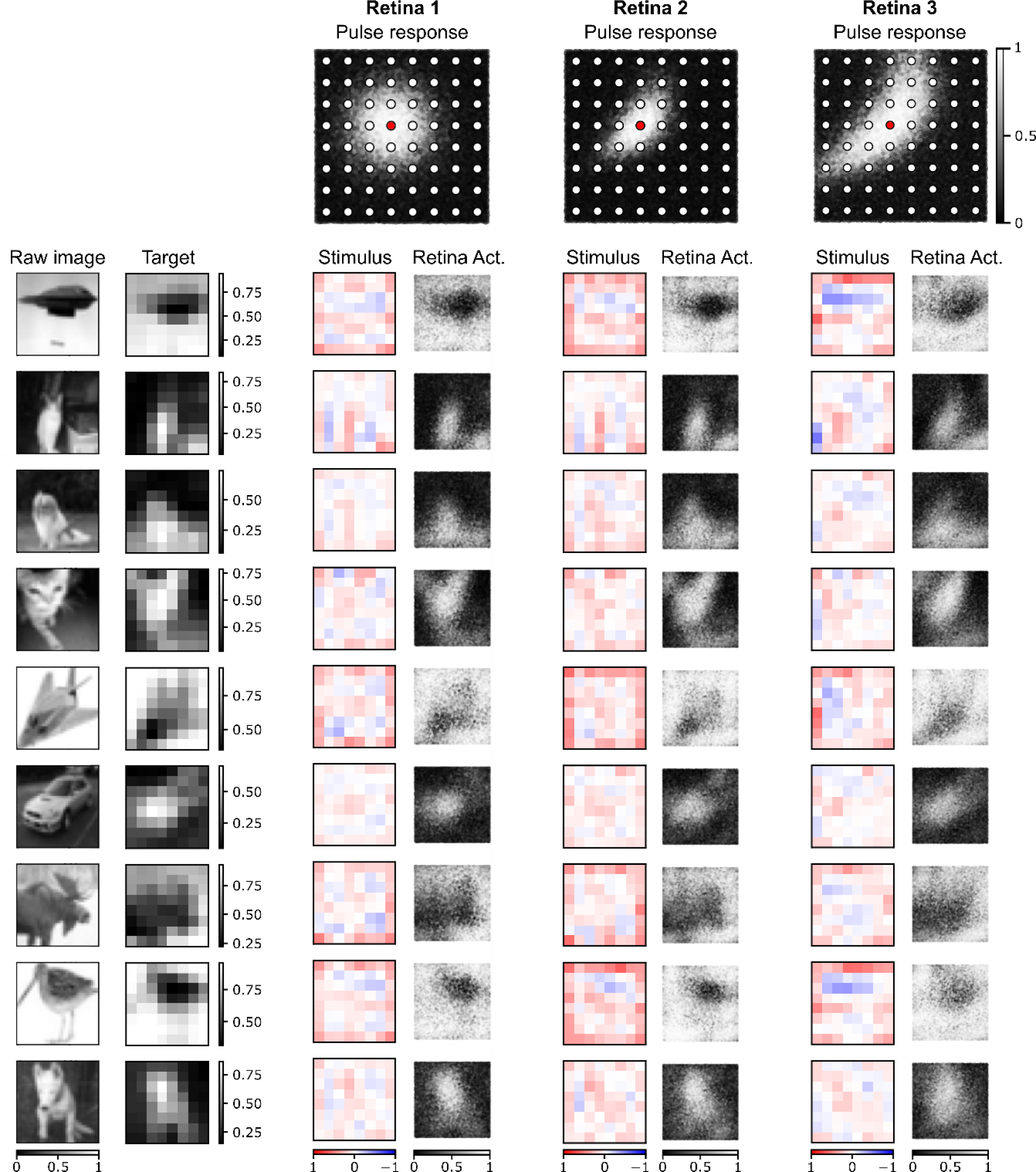
The ANN-based approach produces efficient stimulation patterns in retinas with different response topologies. Qualitative results obtained in three different retinas: Retina 1 – regular topology, identical to those presented in the main paper; Retina 2 – elongated phosphene topology, obtained by changing the ERF parametrizations (y sigma larger than x sigma, ERF angles confined to a smaller range, set as a function of cell coordinates, retina transfer function parameters set as a function of cell coordinates); Retina 3 – elongated phosphene topology, as Retina 2, but with larger spread. The top row shows the pulse response s of each retina to a single pulse with amplitude 1. The remaining rows show the results obtained for different test images. Each retina was trained with a different MPN/SGN. The SGN learns stimulation patterns that adjust to specific retina responses.

**Figure S2.**
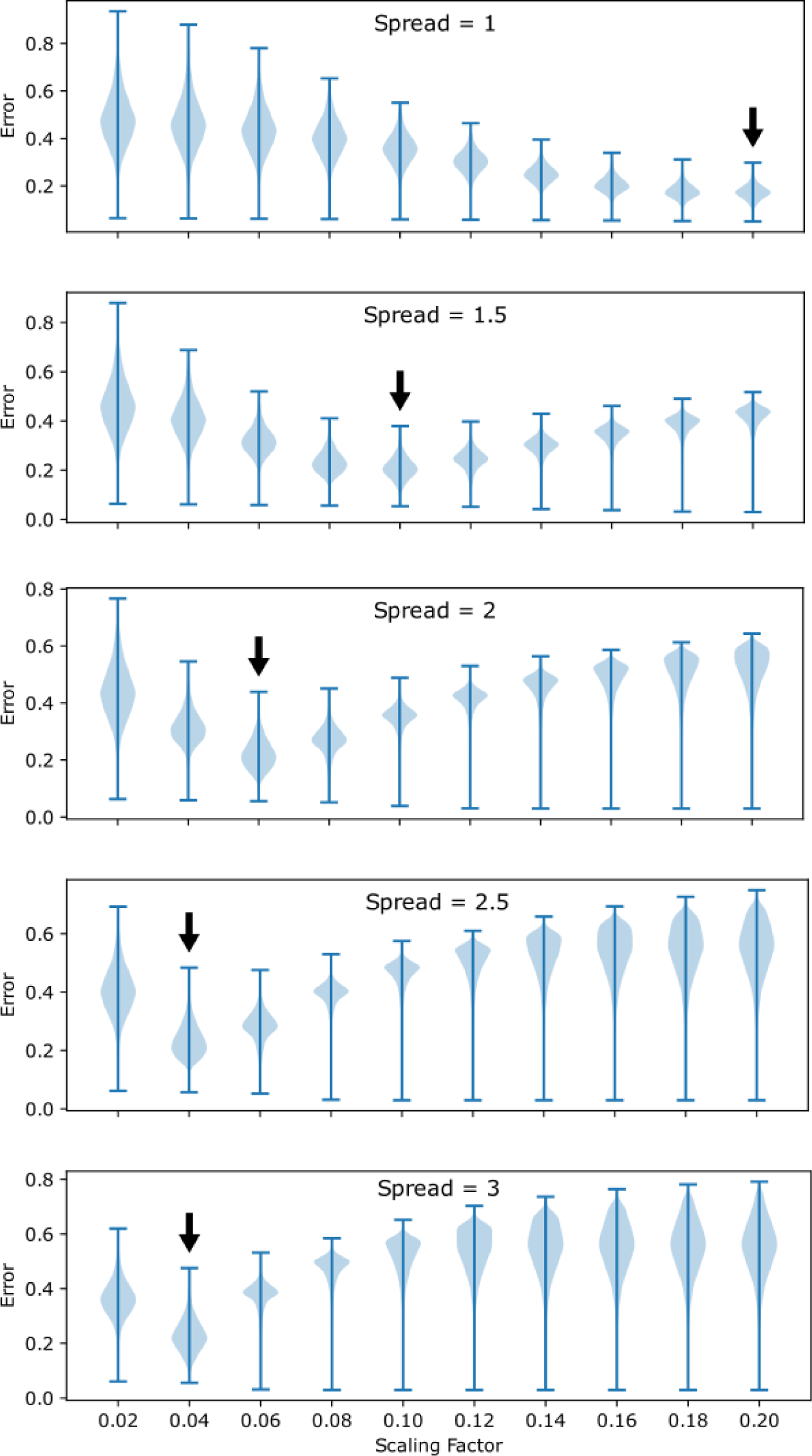
Scaling factor selected for each spread in the conventional stimulation method of. **Figure 8**. To ensure that we compared the ANN approach with the best version of the conventional method, we tested different scaling factors and selected the most efficient one for each spread. We tested each scaling in the 10,000 test images and calculated the RMSE between the retina activation and the measurable target. For each spread, we chose the scaling with lower mean RMSE (black arrow).

**Figure S3.**
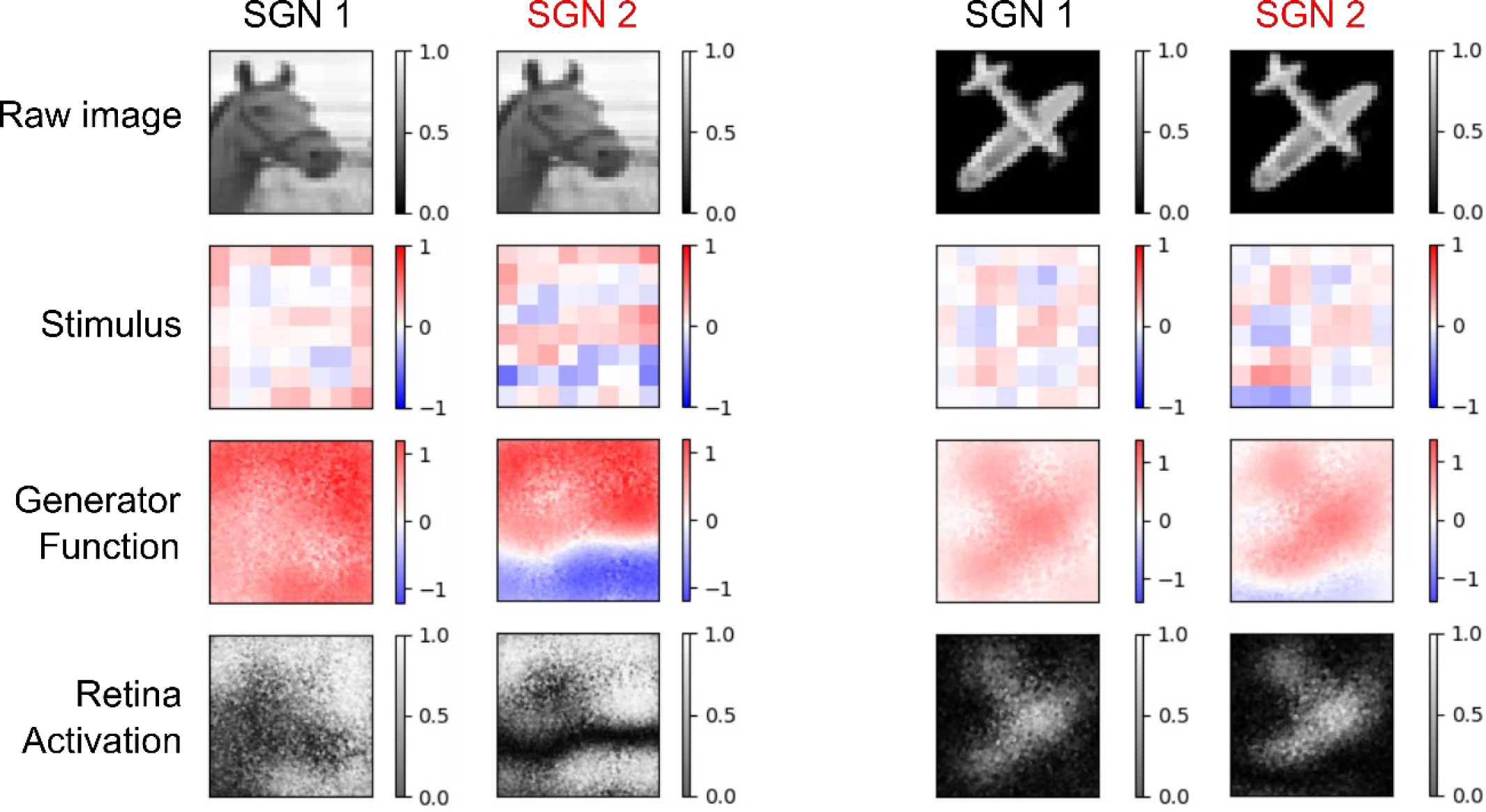
The SGNs that become stuck in local minima during training induce bipolar generator functions 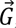 that may lead to artifacts in the retina activation. SGN 1 was the result of a successful training, thus generating unipolar 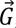 that lead to the expected retina activations. SGN 2 became stuck in a local minimum during training (worst performance in Figure 7.B). The higher training/validation errors obtained with this network are due to the artifact caused by the transition between regions of positive and negative 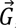. That is, in one region, the retinal cells are being activated due to an overall negative current, whereas in the other region they are being activated due an overall positive current (since the retinal transfer function is a double sigmoid with regards to 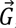, both positive and negative 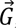 values lead to retinal activation). In the transition between regions, the current is null, and thus there is no retina activation that leads to a dark artifact (example of the left). However, in some situations, this transition region may not affect the results or even be beneficial (example on the right). A network that learns how to effectively exploit these “artifacts” could be used to create sharp regions of no retinal activation, as proposed by Spencer et al., (2021).

